# Imaging and dynamic causal modelling reveal brain-wide changes in effective connectivity and synaptic dynamics during epileptic seizures

**DOI:** 10.1101/160259

**Authors:** RE Rosch, PR Hunter, T Baldeweg, KJ Friston, MP Meyer

## Abstract

Pathophysiological explanations of epilepsy typically focus on either the micro/mesoscale (e.g. excitation-inhibition imbalance), or on the macroscale (e.g. network architecture). Linking abnormalities across spatial scales remains difficult, partly because of technical limitations in measuring neuronal signatures concurrently at the scales involved. Here we use light sheet imaging of the larval zebrafish brain during acute epileptic seizure induced with pentylenetetrazole. Empirically measured spectral changes of spontaneous neuronal activity during the seizure are then modelled using neural mass models, allowing Bayesian inference on changes in effective network connectivity and their underlying synaptic dynamics. This dynamic causal modelling of seizures in the zebrafish brain reveals concurrent changes in synaptic coupling at macro- and mesoscale. Fluctuations of synaptic connection strength and their temporal dynamics are both required to explain observed seizure patterns. These findings challenge a simple excitation-inhibition account of seizures, and highlight changes in synaptic transmission dynamics as a possible seizure generation pathomechanism.

**Abbreviations:**

LFP: local field potential
PTZ: pentylenetetrazole
DCM: dynamic causal modelling
CSD: cross spectral densities
PEB: Parametric Empirical Bayes

## Introduction

Epileptic seizures are characterised by transient disturbances in the brain’s electrical activity that result in changes of patients’ behaviours or perceptions. When recorded through electroencephalography (EEG), the appearance of these paroxysmal electrical discharges often falls within recognisable categories (e.g. spike and wave complexes, paroxysmal fast activity, rhythmic slow waves). Yet even strikingly similar EEG patterns may be caused by different pathological mechanisms ranging from acute exposure to toxins, to genetic mutations as part of a neurodevelopmental syndrome (Shorvon, 2011). Understanding the link between the wide range of *particular* neurological disturbances and the small set of *generic* epileptic dynamics visible on EEG, holds the potential for identifying common pathophysiological mechanisms and ultimately develop novel treatment strategies.

One way to identify the effects of particular pathologies on neuronal systems is the use of animal models, where the effects of different interventions (such as drug treatments, or genetic mutations) can be evaluated *in vivo* (Depaulis et al., 2015; Parker et al., 2011; Seo and Leitch, 2014). Zebrafish in particular have been of recent interest for epilepsy research because they (i) are a vertebrate model organism with basic neuroanatomic similarities to the mammalian brain, (ii) allow relatively easily the introduction of novel genetic mutations (Dhindsa and Goldstein, 2015), or large-scale screening of different drug interventions (Baraban et al., 2013; Griffin et al., 2017) and (iii) allow a range of functional recordings at different observational scales ranging from single neurons (Kibat et al., 2016) to the entire brain (Ahrens et al., 2013). The recent emergence of light-sheet microscopy as a way to functionally record from the whole larval zebrafish brain at single-neuron resolution offers the potential for detailed insights into both the microcircuitry and whole brain dynamics underlying many neurological conditions that can be replicated in this model (Keller et al., 2014).

Insights into the generic nature of seizure dynamics have largely been derived from computational modelling of EEG dynamics (Jansen and Rit, 1995; Lopes da Silva et al., 1974; Lytton, 2008). Dynamic systems theory reveals that there is a small number of mathematically defined transitions in and out of stable oscillatory states that can replicate the transitions between interictal and seizure-like activity (Breakspear, 2005; Izkhikevich, 2000; Robinson et al., 2002). These transitions, or bifurcations, describe mathematically how a system can suddenly change its output from one type of oscillation to another, even with very small changes in the system’s setup. Using simple models of neuronal populations and exploring effects of different model parameters on the occurrence of bifurcations (Jirsa et al., 2014) is a powerful method to identify the relationship between particular neurobiological changes (i.e. parameter changes in the model), and observable features (i.e. bifurcations in oscillatory patterns produced by the model).

Combining the power of *in vivo* models of epileptic seizures (in light of available whole-brain functional imaging techniques) and *in silico* models of seizure dynamics has the potential to lead to an in-depth understanding of how specific disruptions at the microscale lead to whole brain phenotypes recognisable as epilepsy. One strategy to combine computational modelling with novel imaging techniques is the use of dynamic causal modelling (DCM, Friston et al., 2003). Here, a Bayesian model inversion approach is used to fit neuronal models to existing data, allowing inference on which of a number of possible models best explains the empirical observations (Friston et al., 2007; Penny et al., 2010; Stephan et al., 2010). This approach has been successfully applied to a range of different imaging techniques including EEG, magnetoencephalography (MEG), functional magnetic resonance imaging (fMRI), functional near-infrared spectroscopy (fNIRS) and local field potential (LFP) recordings (Daunizeau et al., 2011; Kiebel et al., 2008; Moran et al., 2011a; Razi et al., 2014; Tak et al., 2015). DCM has been applied to test mechanisms underlying seizure generation and spread in scalp EEG (Cooray et al., 2016), invasive recordings in patients (Papadopoulou et al., 2015), and in invasive recordings from *in vivo* animal models (Papadopoulou et al., 2016). However, these approaches remain limited by their limited spatial resolution and resultant difficulty in identifying regionally localised dynamics across a whole-brain network.

**Fig 1.**
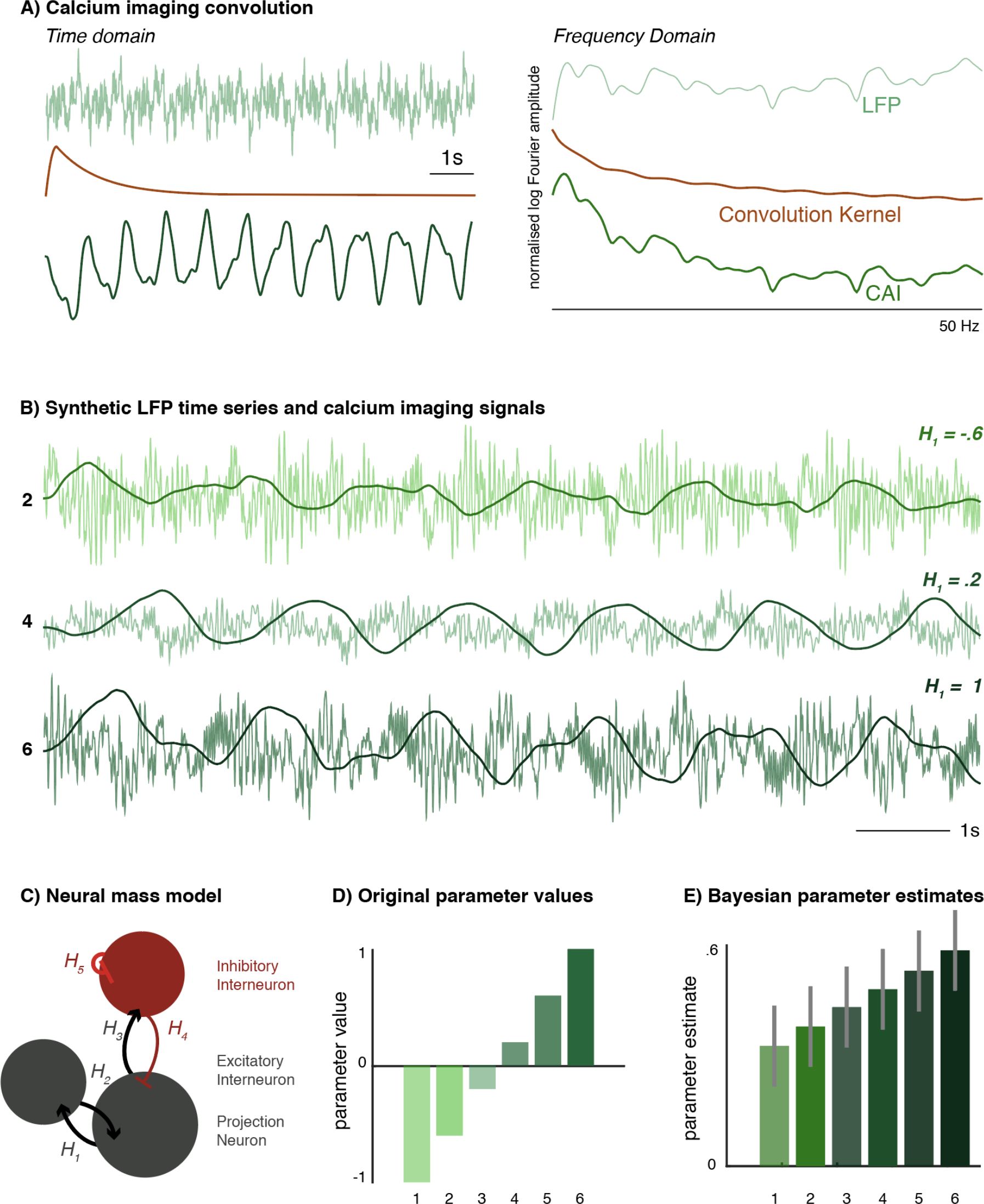
Dynamic causal modelling results of simulated calcium imaging time traces. (A) Calcium imaging dynamics were modelled by convolving LFP-traces (top) with a calcium imaging kernel (middle), resulting in a CAI time trace. The CAI trace follows slow LFP dynamics, whilst attenuating faster components of the original signal. In frequency space (right), the convolution differentially scales low and high frequency components, but preserves most frequency features. (B) LFP-like traces from a three-population neural mass model with increasing values of a single parameter, H_1_- also shown in (*C*). In the time domain, these were then convolved with the CAI kernel, resulting in six different traces (three examples shown). (C) The same neural mass model is subsequently fitted to the CAI traces. Bayesian model comparison (Bayes factor 2.6, not shown) between hierarchical (PEB) model inversion identifies correctly that differences between simulated CAI traces were caused by (D) the effects of variations in the *H*_1_ parameter on the synthetic LFP traces. (E) The DCM approach identifies the increase of *H*_1_ across the six model inversions from the CAI traces, shown here with a Bayesian 95% confidence interval. Whilst the group mean parameter value and the effect size are different, this inversion correctly identifies the linear increase in the parameter from the simulated CAI dataset. *LFP* – local field potential, *CAI* – calcium imaging, *DCM* – dynamic causal model, *PEB* – parametric empirical Bayes

Here we present a novel application of DCM on calcium imaging data from the larval zebrafish brain during epileptic seizures. In this acute chemoconvulsant model, seizures were induced with pentylenetetrazole (PTZ) in healthy larval zebrafish and seizures recorded using brain wide expression of the genetically encoded reporter of neural activity (GCaMP6F) and light sheet imaging across a single slice capturing five main bilateral brain regions. The DCM analysis rests on the notion that these acute seizures are associated both with changes in localised microcircuit dynamics that result in a phase transition between resting activity and the pathological seizure state (Breakspear, 2005; Jirsa et al., 2014), and with measureable changes in whole-brain connectivity (Nehlig, 1998; Omidvarnia et al., 2017; Sinha et al., 2016). DCM allows concurrent testing of the following emerging hypotheses across these different spatial scales: (1) seizures lead to a measureable reorganisation of effective connectivity between regions (Burns et al., 2014), (2) local excitation-inhibition imbalance explains associated regional spectral changes (Netoff et al., 2004), (3) in addition to changes in connection strengths, seizures are also associated with changes in synaptic transmission dynamics (Papadopoulou et al., 2016).

These hypotheses are tested using regional averages of the light-sheet imaging traces and estimating their spectral composition (i.e. cross spectral density) changes over time using a sliding window approach (Papadopoulou et al., 2016) – DCMs are fitted to each of the separate time windows, and hierarchical modelling using parametric empirical Bayes (PEB) (Friston et al., 2015) is used to identify slow fluctuations in synaptic (model) parameters induced by PTZ. The analysis described below comprises the following components: (a) assessment of construct validity of the model using synthetically generated data, (b) identification of an appropriate network architecture using Bayesian model comparison of DCMs estimated for the baseline, pre-seizure data, (c) applying PEB to successive data epochs to identify synaptic changes underlying PTZ-induced seizure activity and test the hypotheses above, (d) using the specified electromagnetic models to describe the transitions through parameter space the zebrafish brain undergoes during a seizure. Our findings suggest that changes in both synaptic connection strength and their dynamics underlie the generation of seizures, challenging the notion that seizures arise from a simple disruption in excitation-inhibition balance.

## Results

### Simulations

In the analysis presented here, we used electromagnetic neuronal mass models originally designed to explain data features observed in LFP recordings. First, we confirmed the construct validity of this approach – i.e. applying DCM for local field potentials to time traces derived from light sheet imaging – by applying the analysis to synthetic data. These were derived from a neural mass undergoing predefined parameter changes: Using a single ‘source’ consisting of three coupled neuronal populations, we generate noisy LFP-like data. These are then convolved with a composite exponential decay kernel modelling calcium probe dynamics (Chen et al., 2013). These surrogate fluorescence time traces are then downsampled to the sampling frequency achieved in the single-slice light sheet imaging (20Hz). This linear convolution equates to a simple addition of the signals in (log) frequency space. Because of the simple frequency composition of the calcium imaging kernel, this linear transformation preserves much of the spectral features in the underlying LFP like signal (Fig 1A).

**Fig 2.**
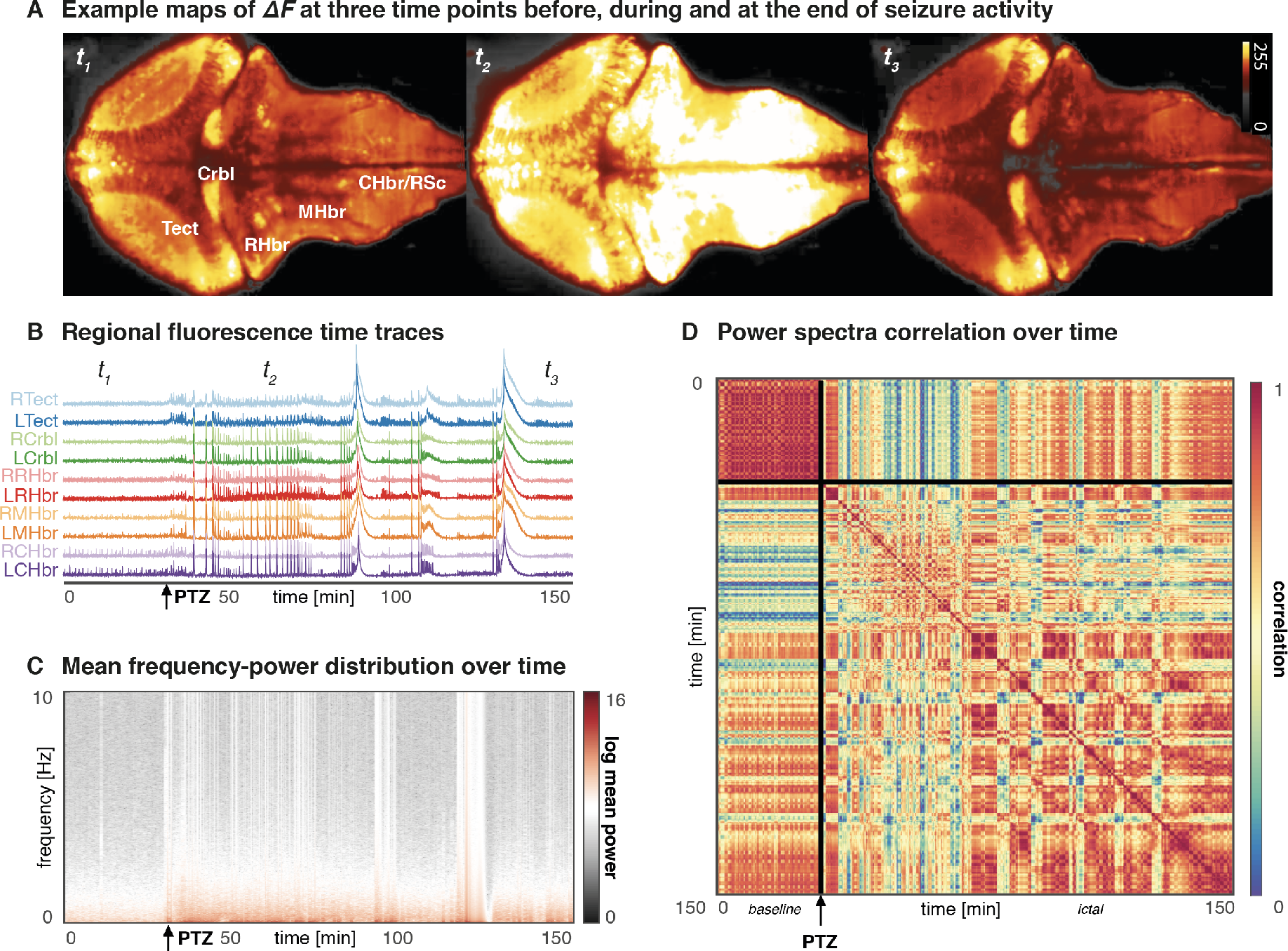
PTZ-induced seizures recorded in the zebrafish larvae using light sheet imaging. (A) This image shows heat maps of fluorescence in a single slice of the intact larval zebrafish brain. Seizure activity (t_2_) is visually apparent as an overall increase in synchronous neuronal activity compared to baseline state (t_1_). (B) An average of the fluorescence signal across 5 bilateral anatomically defined regions disclose seizures as an inrease in generalised high amplitude activity. (C) Time frequency analysis using Fourier transforms on sliding window data segemnts shows that PTZ causes an increase largely in low frequencies (<2Hz), with intermittent bursts of more broadband activity (the figure shown is an average over *n=3* animals). (D) A correlation matrix showing correlation indices of the power-distribution patterns across different time points (delay-delay matrix). This reveals three distinct time periods, corresponding to baseline (<30min), ictal (30-70min) and late ictal (>70min) phases with distinct spectral signatures and temporal dynamics. Tect - Tectum, Crbl - Cerebellum, RHbr - Rostral Hindbrain, MHbr - Mid-Hindbrain, CHbr/RSc - Caudal Hindbrain/Rostral Spinal Cord

The variations in the single neural mass model parameter introduces spectral changes in both the surrogate LFP and fluorescence time traces (Fig 1B). We now fitted a three-population neural mass model (of the kind used to generate the LFP traces, Fig 1C) separately to each of the fluorescence time traces. This yielded six separate dynamic causal models (DCMs), one each fitted to the six time series generated using with variations in a single parameter as shown in Fig 1D. Using a hierarchical parametric empirical Bayesian model, we then identified which parameter could best explain the differences in these DCMs (fitted to fluorescence signals). This successfully identified variations in the correct parameter (*H_1_*) as the most likely cause for the differences in time series. Furthermore, the estimated between-DCM differences in *H_1_* values also captures the direction of the linear change introduced in the original LFP time series.

**Fig 3.**
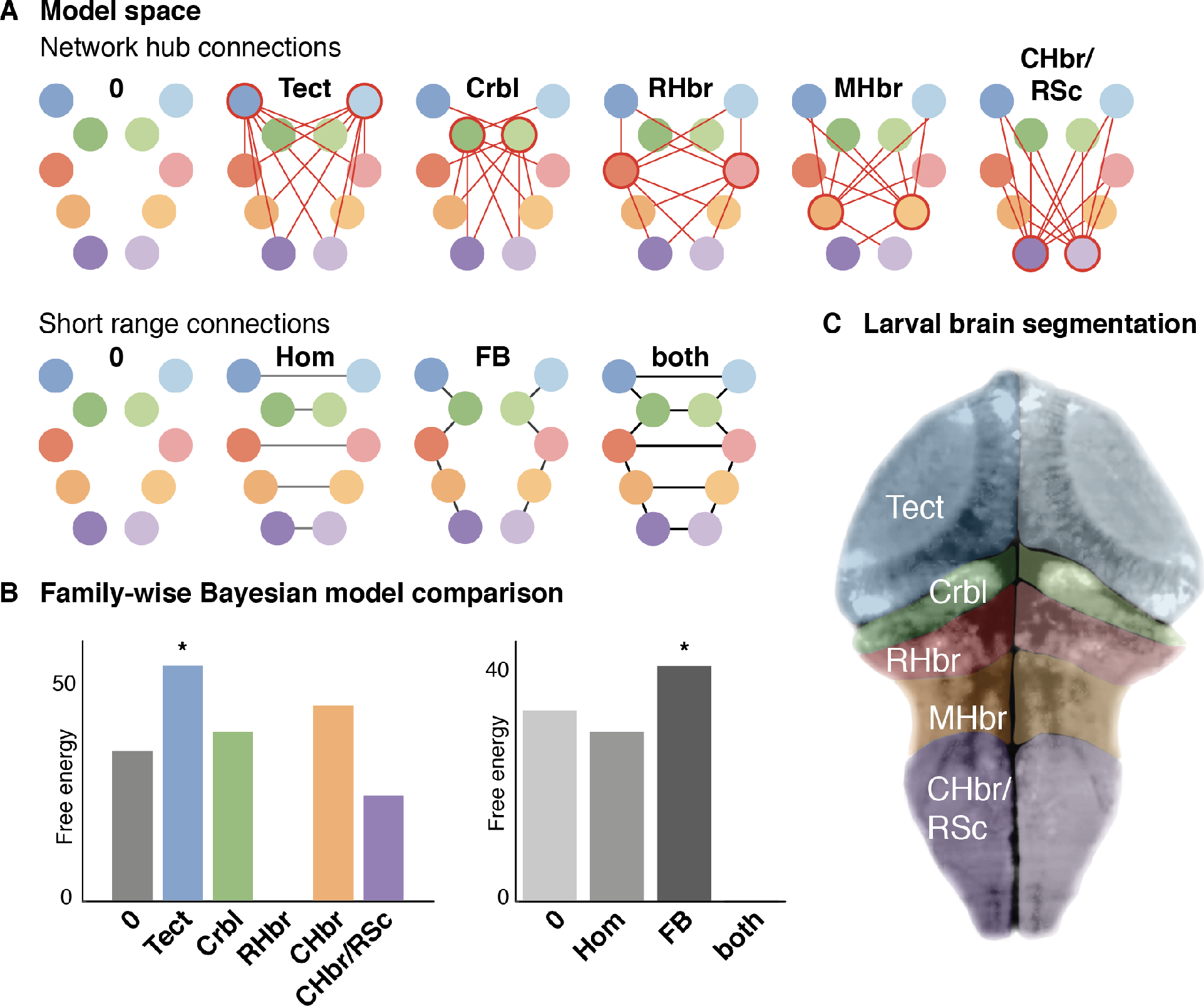
Model comparison of network architecture and first level model predictions. (A) DCMs were inverted for cross-spectral density summaries of baseline data. Bayesian model reduction was used to estimate the model evidence across a number of reduced models that were characterised by the presence or absence of sets of between-region reciprocal connections (neighbouring, homotopic, and hub connections). Each region was modelled as a set of three coupled neural masses including projection neuronal populations, and locally connected excitatory and inhibitory interneurons (B) Bayesian model selection identifies the model with neighbouring, and homotopic connections as well as the optic tectum with hub-like connectivity as the best explanation for the observed dynamics at baseline. (C) Using this model architecture to fit individual DCMs to the sequence of sliding-window cross spectral density estimates provides excellent model fits capturing the fluctuating spectral composition of the raw calcium imaging trace.

### Seizure recordings

In order to elicit epileptic seizures, PTZ was infused in the bath of *n=3* zebrafish. The resultant seizure activity was recorded with light sheet imaging utilising a genetically encoded calcium sensor (GCaMP6F). Neural activity was were recorded from single slices *in vivo* in agarose immobilised larvae. The activity within the whole imaged slice was readily apparent in the fluorescence images (Fig 2A). We divided the slice into 5 bilateral regions of interest to extract fluorescence time series from the recording. These showing distinctive features consistent with highly correlated epileptic seizure activity (Fig 2B). Using a sliding window (length: *60s,* step: *10s*) we could estimate the time-changing frequency content using a Fourier transform, which demonstrate a particular increase in low frequency power after PTZ infusion (Fig 2C), with additional intermittent bursts of broadband activity seen. Estimating correlations between the regional power-frequency distributions across different time windows reveals apparently distinct phases of PTZ induced seizures (Fig 2D): A baseline that is stable over time (0-30 minutes), an initial ictal period that differs most from the baseline state (30-70 minutes), and a late ictal period where time periods of apparent similarity (i.e. high correlation) to the baseline are interrupted by intermittent different (i.e. low correlation) segments (70 – 150 minutes).

### Functional network architecture at baseline

We employed Bayesian model comparison to identify the effective connectivity network that best explains the baseline data. In brief, baseline activity was modelled as spontaneous activity arising from a coupled network of neuronal sources. Each source is made up of a three-population neuronal microcircuit (excitatory, inhibitory interneuron populations, and additional projection neuron) that is fitted to a cross-spectral density summary of the fluorescence signal at baseline. A single fully connected network was fitted as a single dynamic causal model (DCM) inversion. Using Bayesian model reduction and Bayesian model selection we compared models, where specific sets of between-region reciprocal effective connections were either present or absent. These sets of connections were (1) hub-like connectivity between a specific region and all other regions; (2) short range connection between neighbouring, and homotopic brain regions (Fig 3A). Bayesian model comparison across the reduced models in this model space provided evidence that the baseline configuration can best be described as a network of neighbouring connected nodes with the tectum acting as a network-wide hub (Fig 3B).

### Hierarchical dynamic model of seizure activity

Using this model architecture, individual DCMs are fitted to the sequence of sliding-window derived cross-spectral density summaries of the original data. At this stage (i.e. first level models), each time window is modelled as independent DCM. The model fits show that these independently inverted models recreate the dynamic fluctuations of spectral composition observed during a seizure very well and thus provide a good representation of the original data features (Fig 4A).

Parametric empirical Bayes (PEB) can be employed to identify parameters across individual DCMs that vary systematically with specified experimental variables. In brief, PEB allows one to invert hierarchical models where, in this instance, the first level of the model corresponds to a sequence of time windows. The second level of the model then uses the posterior densities over the first level parameters to model changes (here fluctuations) in the first level parameters. We modelled PTZ induced changes as a mixture of four effects: (1) a simple model of PTZ bioavailability as first order pharmacokinetics with a maximum effect achieved at 30 minutes, (2) a tonic effect switched on for the duration of PTZ exposure, (3) a monotonically increasing effect representing the influence of prolonged seizure activity, (4) oscillatory effects at different slow frequencies represented by a set of discrete cosine transforms (Cooray et al., 2015). This approach provides a single model at the group level (i.e. across all time windows, and all individual fish) and parameter changes are modelled as a mixture of experimental and random effects. These second level inversions also provide an estimate of the model evidence, so that different models can be tested against each other.

In the first instance, we compared models where only subsets of between region connections were allowed to vary between time points. Bayesian model comparison shows that only changes in the forward connections to the network hub (i.e. bilateral tectum) are required to explain the spectral changes during seizure activity (Fig 4B). Model comparison was also used to test for PTZ induced changes in the intrinsic coupling parameters in individual regions. There was strong evidence for an involvement of all measured brain regions (Fig 4C). The estimated parameter changes induced by PTZ were varied between different brain regions, but overall showed a relative reduction in excitatory time constants (suggesting faster responses), reduction in inhibitory intrinsic connections, and a reduction the influence of other regions on the optic tectum (i.e. a reduction in forward connections) (Fig 4D).

**Fig 4.**
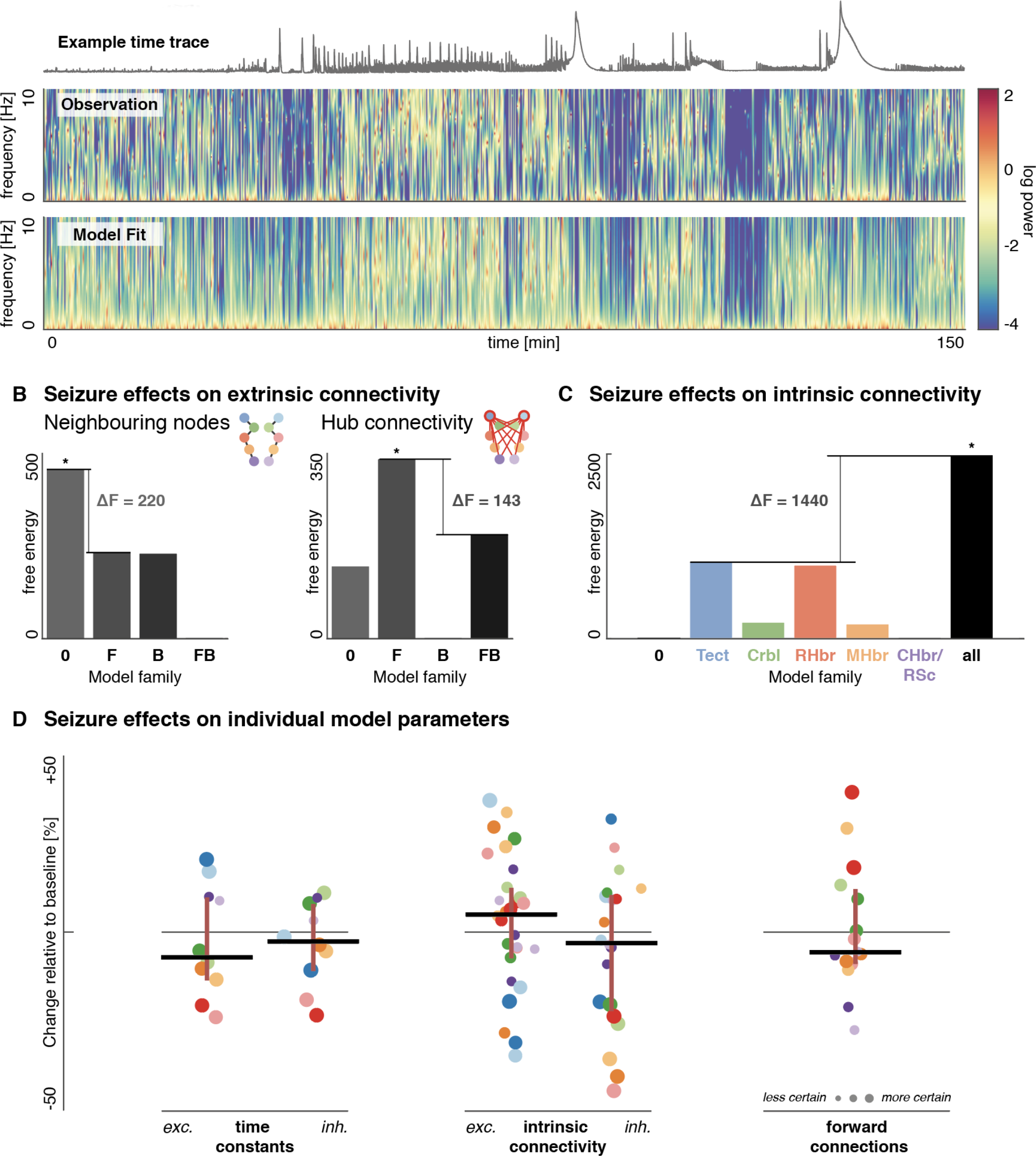
Synaptic coupling changes associated with PTZ-induced seizures. (A) DCMs fitted to individual time windows capture the spectral changes measured during seizure activity. Shown is a frequency-power plot of a single eigenmode of cross-spectral density summaries fluorescence time series of a single animal for the duration of the experiment. (B) Bayesian model comparison across several model families at the second (between time-window) level was performed to compare reduced models with a limited set of fluctuating parameters. For extrinsic (between-region) connections, we compared models with seizure-induced changes in *F* forward, *B* backward, *FB* both, *0* or neither connections. Only changes in connections from other brain regions to the hub region showed evidence of being modulated by the seizure activity. (C) We compared models with intrinsic connectivity changes in none of the brain regions, single brain regions, or all brain regions. There was strong evidence for intrinsic connection changes in all brain regions. (D) Individual parameter changes induced by PTZ varied between brain regions. The dot plots show individual parameter estimates, colour-coded by region with the size encoding the certainty of the estimate (i.e. the inverse covariance). Lines indicate the median, with whiskers showing 25^th^ and 75^th^ centiles respectively.

To further explore the relationship between specific parameter changes and the spectral output, we simulated the spectral output of a single three-population source for a range of different parameter values informed by the PEB analysis above. We extracted the parameter estimates for time constants and intrinsic connectivity within the right tectum (Fig 5A) over time across all components of the PEB model (i.e. tonic seizure effects, monophasic PTZ effect, prolonged seizure effect, discrete cosine transforms, random between-subject effects). We then extracted the first principal component of the intrinsic connectivity changes, and the time constant changes over time (Fig 5B).

**Fig 5.**
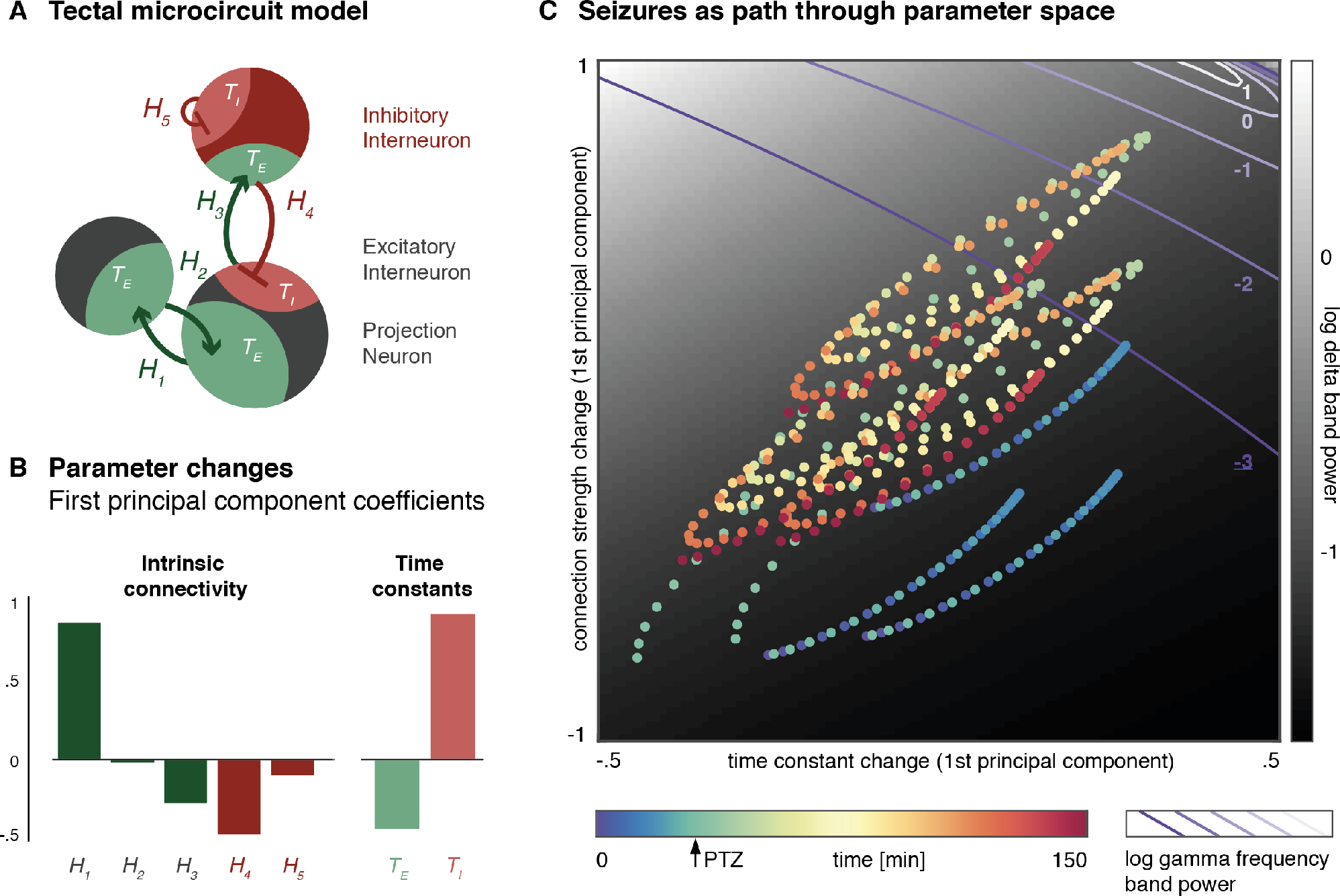
Seizures unfold as path through parameter space. (A) In this simulation a single source consisting of three population is analysed along variations in seven parameters: 5 intrinsic connectivity parameters, and 2 time constants. (B) The coefficients for the first principal component of intrinsic connections (left) and time constants (right) are shown. (C) Individual time windows are plotted onto a two-dimensional parameter space consisting of the 1^st^ component of time constant changes, and the first component of connection strength changes. For each point in this parameter space, simulations of a single source with that particular parameterisation yield an estimate of predicted delta- (black and white heat map) and gamma-power (purple isoclines) respectively.

Plotting each time window onto this reduced parameter space containing most of the variance in the coupling parameters represents the seizures as a spiral path through parameter space. We can apply the parameter combinations at each point in the parameter space to a microcircuit model and predict the spectral output. Here we show log delta band power as a heat map, with log gamma band power superimposed as isoclines (Fig 5C). This forward modelling approach shows that during the seizure, the model enters a section of parameter space characterised by both high delta and gamma power components, which is also seen in LFP recordings during seizures in zebrafish reported in previous studies (Baraban et al., 2013).

## Discussion

In this report we combine light-sheet imaging of the intact larval zebrafish brain during PTZ-induced seizures with dynamic causal modelling in order to identify network-wide connectivity changes induced by the pharmacological intervention. Even well studied pharmacological interventions, such as PTZ show multi-scale effects across the nervous system (Baraban et al., 2005; Huang et al., 2001; Kalueff, 2007; Nehlig, 1998). Thus linking membrane-level changes with the whole-brain seizure phenotype remains challenging. The approach illustrated here allows the comparison of different mechanistic hypotheses and identification of neuronal changes caused by PTZ.

### Validity of DCM for calcium imaging traces of seizure activity

Time series derived from calcium imaging have previously been demonstrated to be highly correlated with underlying LFP changes (Chan et al., 2015). Whilst LFP generally allows measuring of neuronal population activity at a higher temporal resolution (including activity >100Hz), the temporal resolution of the calcium imaging approaches is more limited due to both the sampling frequency (Keller et al., 2014), and the fluorescence decay dynamics of the calcium-sensitive probe (Chen et al., 2013).

The predominant frequency components of both resting brain activity and seizure activity in the larval zebrafish brain are in the delta (<4Hz) and theta (4-8Hz) band (Afrikanova et al., 2013). LFP fluctuations in these frequency bands are largely preserved in calcium imaging, and apparent even at sampling frequencies as low as 20Hz. This is in keeping with the obvious signatures of PTZ-induced seizures both in the raw calcium imaging data, the regional averages, and the frequency domain. Modelling of changes in these low frequency ranges using electromagnetic neuronal models designed for LFPs may therefore be appropriate. Here we have further confirmed this by our construct validity testing using synthetic data: Even after convolving a synthetic LFP-like trace (generated with known parameter values) with a calcium imaging kernel, adding noise, and down sampling to a lower sampling frequency, the DCM analysis correctly identifies the originally manipulated parameter.

This approach does not fully exploit the spatial resolution offered by the calcium imaging data, which will need to be addressed with custom approaches to modelling of individual neurons (Rahmati et al., 2016). However there are specific advantages to using the regionally averaged calcium traces: The average calcium imaging trace is spatially less biased than an LFP trace - whilst LFP recordings pick up on population signal, they are heavily biased towards signals close to the recording site. Light-sheet imaging samples the entire slice in a spatially unbiased fashion and provides a conceptually closer approximation to the assumptions made for the neural mass models used in DCM (Moran et al., 2013). Heuristically, this spatial averaging suppresses local fluctuations very much in the same way that averaging over time in the event related potential studies (in electrophysiology) reveals dynamics that are conserved over multiple realisations. Furthermore epileptic seizures are an emergent property at the level of neuronal populations, and computational models specifically addressing this ‘mesoscale’ may yield important insights about emergent population-wide features that is less readily apparent from microscale modelling of individual neurons (Kuhlmann et al., 2015).

### Network organisation in the larval zebrafish brain

DCM allows for the estimation of network coupling parameters underlying neurophysiological recordings. Early during zebrafish development, retino-tectal connections develop and stereotyped but effective visuomotor behaviour is established (Meyer, 2006; Niell and Smith, 2005; Niell et al., 2004; Portugues and Engert, 2009). This is associated with distributed network activity involving information flow from the optic tectum to other brain areas. This visually-dominated early network activity is also apparent in the DCM analysis, where the tectum has been identified as a hub with widespread connectivity to the rest of the larval zebrafish brain from resting state light sheet recordings at baseline.

This network organisation is modulated during seizure activity, where the modelling approach identifies a reduction of the effective forward connections from other brain areas to the optic tectum. This asymmetric shift in connectivity (with only forward, but not backward connections affected), may be indicative of a key role of the optic tectum - as a central network hub at baseline - in driving network-wide synchronisations during an epileptic seizure. The selective reduction in effective connectivity corresponds to previously reported seizure-related changes in functional connectivity estimated from human EEG recordings, where increased clustering during a seizure has been described (Schindler et al., 2008).

Fluctuations in effective connectivity between regions is usually thought of as resulting from changes in direct synaptic connectivity (Nam et al., 2004). However, the asymmetric involvement of a single brain region – where only effective connectivity *to* (and not *from*) the optic tectum is reduced – suggests that local microcircuitry changes may underlie the macroscale changes. This phenomenon has been formally described in other modelling work through a slow local permittivity variable that governs synchronisation between different brain regions and represents different slowly unfolding changes in local energy and metabolic milieu (Proix et al., 2014). The relationship between local and macroscale network changes in epilepsy in the context of hierarchically coupled brain areas is discussed elsewhere (Omidvarnia et al., 2017). In our approach here, slow local changes may appear as the sort of changes in directed effective connectivity estimated for the optic tectum, linking local microcircuitry abnormality with pathological brain-wide reorganisation during the epileptic seizure.

### Intrinsic coupling changes disrupt excitation-inhibition balance in the temporal domain

PTZ acts as an acute chemoconvulsant in a range of different model organisms, likely due to allosteric inhibition of GABA-A receptors (Huang et al., 2001). Previous work on a PTZ rat model showed dose-dependent regionally specific cellular activation (Nehlig, 1998), suggesting differential susceptibility of different brain regions to PTZ effects. Bayesian model comparison of seizures recorded from the zebrafish in this report indicate that changes of intrinsic neuronal population coupling were required in each of the brain region to explain the observed seizure patterns, suggesting that direct effects of PTZ changes in all brain regions are required, rather than pathological changes in a single brain region driving all of the observed seizure effects.

These effects varied widely between different brain regions. This in part reflects different baseline configurations of the regional source models, which in turn require different shifts in coupling parameters in order to achieve the sort of spectral output observed across all brain regions during the seizure. However, overall, the PTZ-related changes are broadly consistent with our current understanding of PTZ effects at the neuronal membrane. Specifically, PTZ is expected to cause a relative decrease of inhibitory connectivity compared to excitatory connectivity; and preferential blockade of fast GABA-A (and not GABA-B) mediated transmission would be expected to cause an increase in the relative inhibitory transmission time constants (i.e. slowing down), compared to excitatory synaptic dynamics – both of these effects are observed in the parameters estimated across the whole brain slice here. Changes in these population-level time constants may relate to the relative contributions of GABA-A and GABA-B transmission, changes in single channel kinetics induced by PTZ (Huang et al., 2001), but are also affected by current input load (Koch et al., 1996), and within-population recurrent connections that are not otherwise explicitly modelled in the neural mass approach presented here (Chaudhuri et al., 2014).

Further exploration of individual parameter effects at a single brain region supports the notion that seizure dynamics in this recording are largely caused by two main effects: a relative disturbance in excitation / inhibition balance with increased excitation and decreased inhibition, and a reciprocal disturbance in the dynamics of excitatory and inhibitory connectivity with slower inhibition and faster excitation. Because we are have fitted fully generative neural mass models, we can make predictions about the spectral output caused by particular parameter combinations beyond the measured ≤10Hz frequency range. This approach reveals that particularly the time points where both connectivity and time effects changes reach their respective extremes, the typical seizure spectral output containing high amplitudes in both low (i.e. delta) and high (i.e. gamma) frequency components emerges.

Recurrent neuronal loops with a close balance of overall excitation and inhibition underlie spontaneous brain activity. The brain is believed to operate near a transitional state from which both subcritical, random dynamics and supercritical, ordered dynamics can emerge (i.e. self-organised criticality, cf. Rubinov et al., 2011). Blocking of the largely GABA-A mediated local recurrent inhibition shifts this balance and allows ordered, seizure-like activity to occur (Shu et al., 2003). In our model the emergence of seizure dynamics requires changes in both connection strengths and their temporal dynamics.

## Conclusion

The analysis presented here illustrates the use of computational modelling to explain neuronal dynamics in the larval zebrafish brain during acutely induced seizures. This approach exploits the spatial independence of light-sheet recordings of brain regions and uses dynamic causal modelling to identify the mechanisms underlying seizure dynamics. This approach allows translating observations from whole-network novel light sheet imaging to the concepts and models used to explain electrophysiological abnormalities observed during seizures.

Seizures in this model are associated with an asymmetric decoupling of the network hub, and changes in excitation/inhibition balance that crucially also involve the temporal dynamics of excitatory and inhibitory synaptic transmission. Mapping the expected spectral changes along both the connection strength and time constant domains of the model within the pathophysiological range estimated from acute seizures allows us the independent contribution of changes in either type of parameter to the overall dynamics. This is the first step to establishing network-wide mechanisms that underlie seizures and may be targeted with novel treatments for epilepsy.

## Methods

### Contact for Reagent and Resource Sharing

Please contact R.E.R. for reagents and resources generated in this study.

### Experimental Model and Subject Details

#### Zebrafish Maintenance

Zebrafish were maintained at 28.5°C on a 14 h ON/10 h OFF light cycle. Transgenic line used: Tg(elavl3b:GCaMP6F) (Dunn et al., 2016). This work was approved by the local Animal Care and Use Committee (King’s College London) and was performed in accordance with the Animals Experimental Procedures Act, 1986, under license from the United Kingdom Home Office.

### Method Details

#### Construction of Light-sheet microscope

The light-sheet design was based on that described in (Wolf et al., 2015). Briefly, excitation was provided by a 488nm laser (488 OBIS, Coherent) which was scanned over 800*μ*m in the Y direction by a galvanometer mirror (6215H/8315K, Cambridge Technology) creating an illumination sheet in the XY-plane. The sheet was associated with two pairs of scan and tube lenses, scanned along the z-axis using a second galvanometer mirror (6215H/8315K, Cambridge Technology) and focused onto the specimen via a low NA illumination objective (5 × 0.16NA, Zeiss EC Plan-Neofluar). The detection arm consisted of a water-immersion objective (20 × 1 NA, XLUMPlanFL, Olympus) mounted vertically onto a piezo nanopositioner (Piezosystem Jena MIPOS 500) allowing alignment of the focus plane with the light sheet. The fluorescence light was collected by a tube lens (150 mm focal length, Thorlabs AC254-150-A) and passed through a notch filter (NF488-15, Thorlabs) to eliminate 488 nm photons. The image was formed on a sCMOS sensor (PCO.edge 4.2, PCO). The 20x magnification yielded a field of view of 0.8 × 0.8 mm^2^ with a pixel dimension of 0.39*μ*m^2^. The detection arm and specimen chamber were mounted on two independent XY translation stages to allow precise alignment of the specimen, detection axis and light sheet.

#### Imaging

Nonanesthetized *Tg(elavl3b:GCaMP6F*) larvae, 5 days post fertilisation, were immobilized at in 2.5% low melting point agarose (Sigma-Aldrich) prepared in Danieau solution and mounted dorsal side up on a raised glass platform that was placed in a custom-made Danieau-filled chamber. Pentylenetetrazole (Sigma-Aldrich) was added to the Danieau-filled chamber after 30 minutes of baseline imaging to a final concentration of 20 mM. Functional time-series were acquired at a rate of 20 Hz, 4x4 pixel binning (1.6 *μ*m × 1.6 *μ*m resolution). Time-series were aligned to a mean image of the functional imaging data for each fish (rigid body transformation as implemented in: SPM12, http://www.fil.ion.ucl.ac.uk/spm/software/spm12). Mean fluorescence traces were then extracted from ten anatomically defined regions of interest for further analyses.

### Quantification and Statistical Analysis

#### Estimation of spectral data features

Mean fluorescence traces from the regions of interest were treated as multichannel time series for subsequent analysis. Short segments derived from a sliding window (length: 60s, step size: 10s) were used to estimate time-varying changes in the spectral composition of the time series: For each step of the sliding window the real component of the Fourier spectrum was calculated. A correlation matrix of region-specific mean Fourier amplitude across all time point was used to visualise slow fluctuations in distributed activity (Betzel et al., 2012; Rosch et al., 2017a). Averages of the windowed Fourier spectra and the power correlation matrix across the studied animals are shown.

#### Simulated calcium imaging traces

To test the construct validity of the inversion approach, we used a neural mass model with known parameterisation to generate an LFP output, convolved this output with a calcium-imaging kernel, and inverted a DCM on those synthetic calcium-imaging traces to test whether the original parameterisation can be reconstructed.

The model was a standard prior three-source neural mass model implemented as ‘LFP’ model in the SPM12 model library (Moran et al., 2013). We generated 6 segments of LFP-like model output with linear variation of a single parameter (*H_1_*) from *-1* to *+1*. The convolution kernel was constructed from a fast inverted quadratic rise lasting *t_up_* = 250*ms* of the form: *y* (*t*) = 2*t* * *t_up_* – *t* ^2^. This is followed by an exponential decay function of the form: *y* (*t*) = 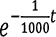. Both functions were normalised so that *y* (*t_up_*) = 1. Parameters of these functions were chosen to approximate the reported dynamics of GCaMP6F (Chen et al., 2013).

#### Inversion of simulated calcium imaging traces

Each individual synthetic calcium imaging trace was inverted using a DCM for cross spectral densities approach with a single three-population neural mass model (Moran et al., 2011b). Using a parametric empirical Bayes approach, we then compared the evidence for models, where changes in a single one of the parameter explain the difference between segments (Friston et al., 2016; Rosch et al., 2017b). Parameter estimates for the winning parameter are then compared to the ‘ground truth’ parameter changes originally introduced into the generative model, therefore providing evidence for which parameter is changed, and how that parameter is changed to achieve the spectral changes contained in the time series.

#### Dynamic causal modelling of empirical calcium imaging traces

<u>Baseline architecture</u>: To characterise functional network architecture at rest, in an initial step only baseline data were analysed using a DCM approach. Specifically, 4-minute segments prior to PTZ were inverted using a single fully connected DCM containing 10 standard prior ‘LFP’ type sources (Moran et al., 2013). Based on the full model inversion, smaller subsets of models were then compared with computational efficiency using Bayesian model reduction, which allows Bayesian model selection for the network architecture that best explains the baseline data (Friston et al., 2016). Model space was designed around three main features: the presence or absence of hierarchical connections between neighbouring brain regions; the presence or absence of homologous connections between bilateral brain regions; and the presence of absence of hub-like connections from one set of brain regions to all other regions (Fig 3).

<u>Seizure data inversion</u>: Based on the dynamic network architecture identified in the step above, an additional DCM analysis was performed to identify slow fluctuations of synaptic parameters within this architecture that could explain seizure activity. For this, data were again divided into segments using a sliding window approach (60s, 50s steps) for each animal separately. DCMs with the architecture derived from the step above were inverted separately for each individual time window.

We then constructed a second level model to estimate between-time window variations in parameters using a parametric empirical Bayesian approach (Friston et al., 2016; Papadopoulou et al., 2016). This contained several temporal basis functions that in combination can explain a majority of possible parameter trajectories: (1) an ‘on/off’ tonic seizure effect step function with onset at PTZ injection; (2) a monophasic seizure effect function with onset at PTZ injection; (3) a linear increase with onset at PTZ injection; (4) a set of three discrete cosine basis functions to model; (5) a set of three regressors modelling random between-fish effects.

This approach provides estimates for how between-time window parameter changes can be modelled as a linear combination of the basis sets provided, as well as a free energy estimate for the model evidence. We can thus perform Bayesian model reduction and selection at this second level, comparing competing model families where only subsets of parameters are free to vary between time windows, and thus select a subset of parameters that best explain the observed changes over time. We broadly divided the model space of these between time-window (i.e. between individual DCM) effects into (a) models with variations in hierarchical coupling, (b) models with variations in hub coupling, and (c) models with variations in intrinsic synaptic coupling parameters as outlined in Fig 4. Family-wise Bayesian model selection was used to select relevant parameters, which were freed in a single model to provide parameter estimates at time window with the estimated maximum PTZ effect.

<u>Forward modelling</u>: To further explore the effects of specific parameter changes, the optic tectum with its hub-like position in the network was analysed further. Parameter changes derived from the linear combination of the temporal basis sets were collated and grouped into time constant and connection strength changes. The first principal component of each of these categories was then used to project the parameter changes into a two dimensional plane. Because the DCM provides a fully parameterised model, we can estimate the predicted spectral output for each point across this plane, by adding the respective principal component values to the baseline parameterisation of the model and simulating its output. We plotted the resultant low frequency (delta-range), and high frequency (gamma-range) power across the parameter space, to indicate how movement in parameter space affects the spectral output.

### Data and Software Availability

#### Software

Analysis in this study was built on tools available as part of the academic freeware package ‘Statistical Parametric Mapping 12’ (www.fil.ion.ucl.ac.uk/spm). This toolbox and all custom code runs on Mathworks© Matlab (https://uk.mathworks.com/products/matlab.html). Custom code is freely available as a github repository (http://github.com/roschkoenig/Zebrafish_Seizure)

#### Data Resources

Extracted time series from manually defined brain regions; windowed data used for DCM analysis (http://github.com/roschkoenig/Zebrafish_Seizure).

